# PopGenAgent: An Agent-Mediated Workflow for Population Genetics Analyses

**DOI:** 10.64898/2026.03.02.709209

**Authors:** Houcheng Su, Weicai Long, Juning Feng, Yusen Hou, Yanlin Zhang

## Abstract

Population genetics analyses typically require orchestrating multiple tools across heterogeneous tasks, from data curation to inference and visualization. In practice, a central bottleneck lies not only in reproducibility, but in maintaining consistency between intermediate diagnostics, analytical decisions, and downstream results as analyses evolve. Here we present PopGenAgent, an agentic workflow that formalizes population-genetic analyses as explicit, multi-step plans with declared inputs, outputs, and dependencies. Each step is instantiated from curated tool and visualization templates, while all intermediate artefacts are preserved within a unified, inspectable context. An interactive agent interface supports iterative plan construction and revision, and leverages accumulated outputs to assist interpretation and report drafting. In a case study of 26 populations from the 1000 Genomes Project, starting from a filtered PLINK dataset, PopGenAgent coordinated analyses including ROH, LD decay, PCA, ADMIXTURE, TreeMix, and *f*_3_. By structuring analyses as explicit, revisable plans with preserved intermediate states, PopGenAgent provides a transparent and extensible environment for conducting population-genetic analyses.

## Introduction

Population genetics studies combine multiple analyses to characterize genetic diversity and population structure Hamilton (2021). In practice, however, the main challenge is often not running an individual program, but keeping file transformations, parameter choices, intermediate diagnostics, and downstream biological interpretation coherent as an analysis is revised. A typical study may move repeatedly across genotype-format conversions, pruning decisions, structure analyses, diversity summaries, and admixture diagnostics before results are ready for reporting.

Reproducible workflow systems alleviate part of this burden by standardizing software environments and execution patterns Wratten et al. (2021). General frameworks such as nf-core Ewels et al. (2020), GenPipes Bourgey et al. (2019) provide reusable pipelines for common analyses, while population genetics specific platforms extend this paradigm to domain workflows: for example, the Pop-Gen Pipeline Platform (PPP) structures VCF processing and downstream utilities Webb et al. (2021), and PAPipe integrates whole-genome processing with predefined population-genetic analyses Park et al. (2024). Despite these advances, a substantial portion of analytical effort remains outside the formalized pipeline structure, arising in the iterative stages where users inspect intermediate outputs, adjust filtering or pruning decisions, and ensure that downstream analyses and summaries remain consistent with those evolving choices Kopelman et al. (2015); Li and Liu (2018).

This issue is particularly acute in population genetics because interpretation usually depends on comparing several complementary outputs rather than on a single endpoint. For instance, analysts often examine heterozygosity and ROH summaries, LD decay, PCA, model-based ancestry inference, and admixture diagnostics together, while also managing conversions among tool-specific formats and plot-ready inputs Kamvar et al. (2015); Leigh et al. (2015). When an upstream choice changes, the practical challenge is not only rerunning commands, but also maintaining a clear record of what changed, which outputs should be regenerated, and how revised results should propagate into figures and narrative text.

Large language models offer a complementary interface for reducing scripting burden and supporting interactive analysis Zhao et al. (2023); Chang et al. (2024); Kasneci et al. (2023). In computational biology, early LLM-based systems have demonstrated tool use and workflow assistance in domains such as genome analysis and transcriptomics Zhou et al. (2024); Huang et al. (2025); Su et al. (2025). For population genetics, however, the main need is not open-ended code generation. It is reliable coordination of standard analyses under realistic failure modes, together with publication-ready outputs that remain traceable to the intermediate artefacts from which they were derived. An effective system therefore needs explicit workflow state, inspectable execution steps, and support for stepwise revision rather than one-shot script generation alone.

PopGenAgent was developed to support this mode of analysis by providing a workflow framework that maintains analysis plans, execution steps, intermediate artefacts, and reportable outputs within a unified context. Analyses begin from prepared genotype datasets and are structured as explicit multi-step plans with declared inputs and expected outputs, instantiated from curated tool and visualization templates. An agent-mediated interface supports iterative plan construction and revision, while leveraging accumulated outputs to assist interpretation and reporting. In the following, we describe the analytical coverage and system design, and illustrate the workflow through a 26-population case study from the 1000 Genomes Project.

## Results

### PopGenAgent represents multi-step analyses as explicit workflows

PopGenAgent separates analytical reasoning from code generation by constraining the agent to a curated set of implemented analysis tools and plotting scripts. Rather than relying on the LLM to generate analysis code dynamically from its own knowledge, PopGenAgent enables the agent to assemble workflows by selecting and parameterizing predefined computational and visualization components. This design improves reliability and analytical consistency by reducing uncontrolled code synthesis, while allowing analyses, intermediate artefacts, and final outputs to remain organized within a persistent workflow context (Fig. 1).

**Figure 1.**
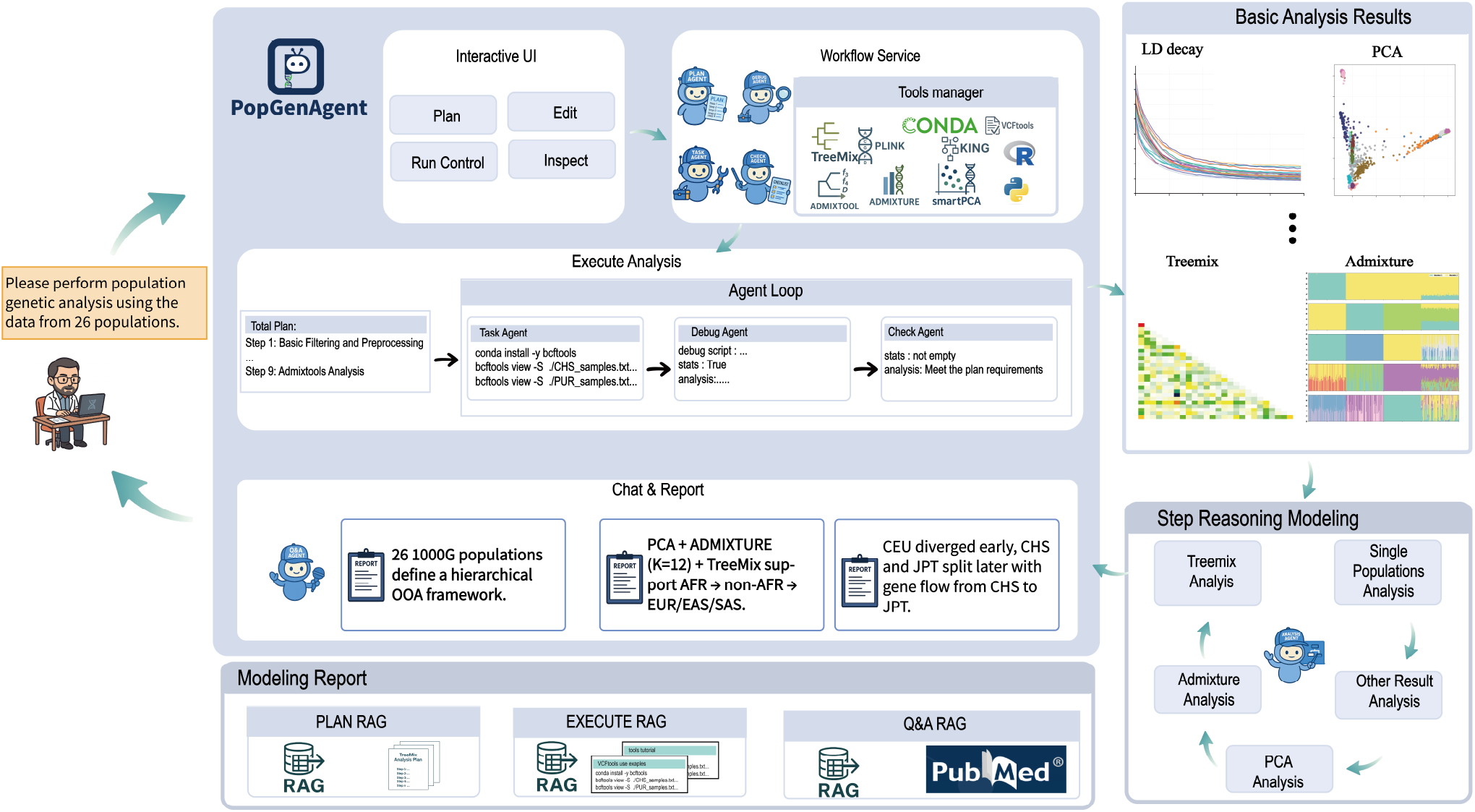
Overview of PopGenAgent. PopGenAgent uses an agent-mediated interface to translate an analysis goal and genotype dataset into an explicit multi-step plan assembled from curated templates. Steps are executed via external analysis tools and plotting scripts, while intermediate and final artefacts are organized for inspection and revision within the same analysis context. The agent can also provide workflow-oriented assistance, including, when configured, literature-backed Q&A and Markdown report drafting grounded in the saved outputs.

PopGenAgent covers tools spanning data preparation and quality control, within-population diversity summaries, population-structure analyses, etc. (Table 1). In the interface, these analyses are represented by plans with declared inputs and expected outputs (Supplementary Fig. S1) and supported by stored tool references together with reusable task and visualization templates (Supplementary Fig. S2). The coverage summarized in Table 1 corresponds to the workflows exercised in the 1000 Genomes example below.

**Table 1.**
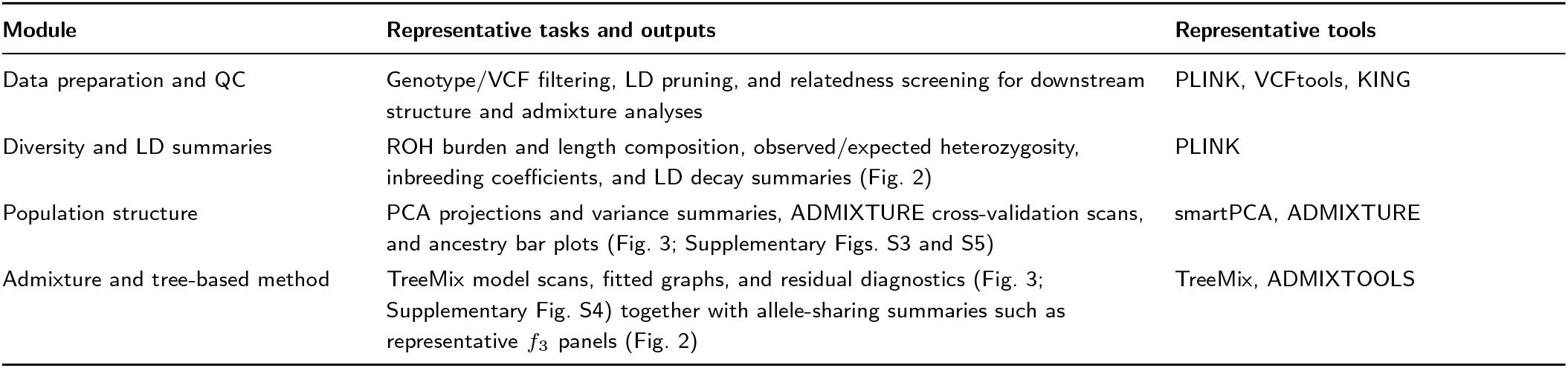
Representative analytical coverage implemented in PopGenAgent and illustrated in this manuscript. The table summarizes core modules, outputs, and external tools coordinated by PopGenAgent.

| Module | Representative tasks and outputs | Representative tools |
| --- | --- | --- |
| Data preparation and QC | Genotype/VCF filtering, LD pruning, and relatedness screening for downstream structure and admixture analyses | PLINK, VCFtools, KING |
| Diversity and LD summaries | ROH burden and length composition, observed/expected heterozygosity, inbreeding coefficients, and LD decay summaries (Fig. 2) | PLINK |
| Population structure | PCA projections and variance summaries, ADMIXTURE cross-validation scans, and ancestry bar plots (Fig. 3; Supplementary Figs. S3 and S5) | smartPCA, ADMIXTURE |
| Admixture and tree-based method | TreeMix model scans, fitted graphs, and residual diagnostics (Fig. 3; Supplementary Fig. S4) together with allele-sharing summaries such as representative $f_3$ panels (Fig. 2) | TreeMix, ADMIXTOOLS |

### Explicit plans and stored artefacts support intermediate-state review

A core feature of PopGenAgent is that workflow intent remains visible throughout an analysis. The saved plan lists step descriptions, required inputs, and expected outputs (Supplementary Fig. S1), making it possible to review what will be executed and which artefacts should be produced before later steps are run. Intermediate plots and tables can then be examined alongside the plan that generated them, and upstream steps can be revised before downstream structure or admixture summaries are regenerated.

To support consistent execution across analyses, PopGenAgent maintains stored tool references and reusable task and visualization templates (Supplementary Fig. S2). These resources capture command patterns, configurable parameters, and expected file organization for supported workflows. Because planning artefacts and generated outputs are preserved within the same analysis context, the agent can also provide output-grounded assistance and assemble report drafts that link narrative text to the saved figures and tables (Fig. 1).

### A 1000 Genomes example illustrates routine downstream outputs from population-genetic analyses

We analyzed the 26-population panel from the 1000 Genomes Project Auton et al. (2015) to illustrate a routine set of downstream outputs generated by PopGenAgent. The analysis started from a filtered PLINK dataset, and derivative files such as LD-pruned inputs and EIGENSTRAT conversions were generated as required by downstream methods. Figures 2 and 3 show representative outputs from this analysis, and Supplementary Figs. S3–S5 provide the corresponding model-scan panels and supporting summaries, including the full ADMIXTURE *K* scan, additional TreeMix fits, and the PCA scree plot.

**Figure 2.**
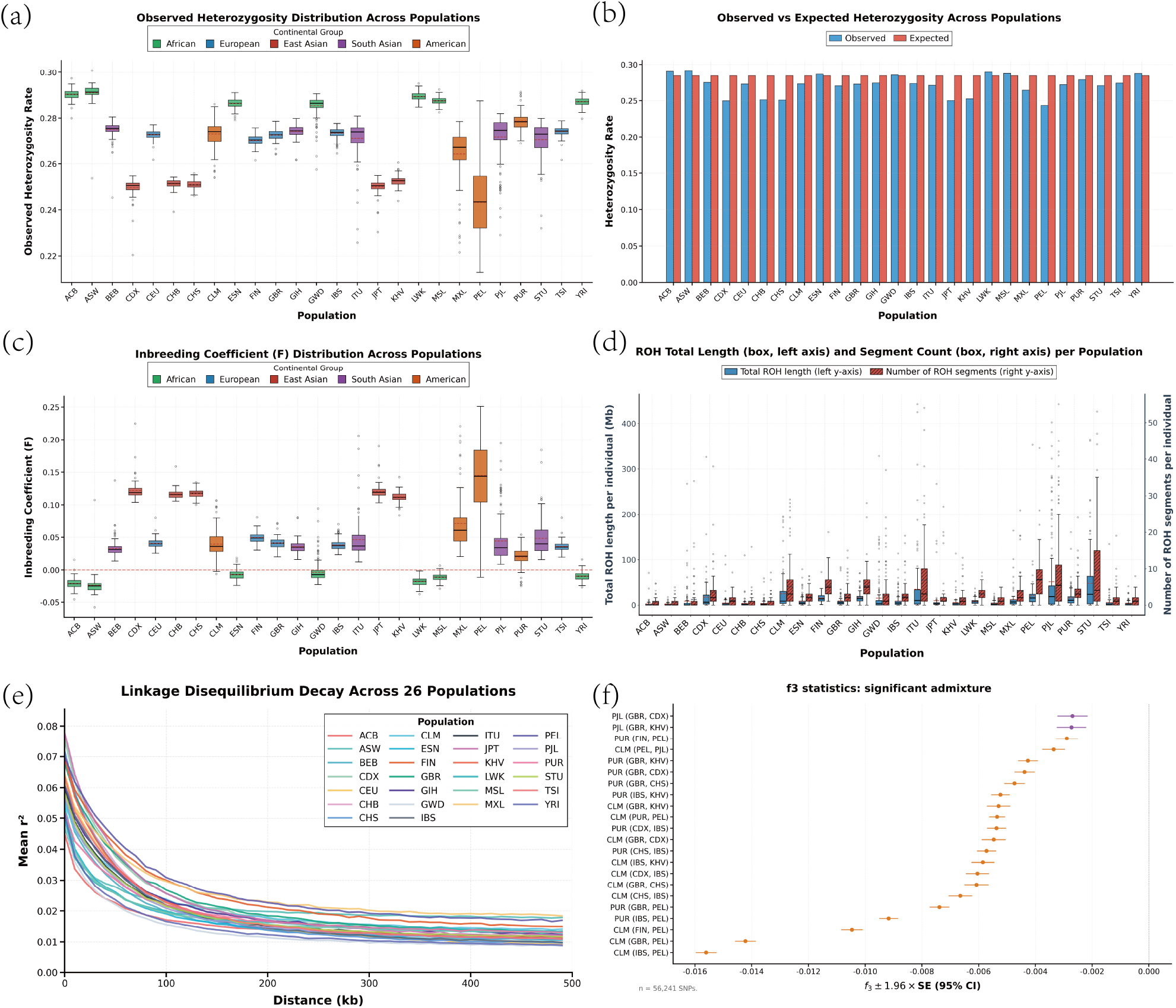
Summary statistics from the 1000 Genomes example analysis. (a) Observed heterozygosity distribution across populations. (b) Observed versus expected heterozygosity across populations. (c) Inbreeding coefficient (*F*) distribution across populations. (d) Distribution of total ROH length and ROH segment count across populations. (e) Genome-wide LD decay curves across populations. (f) Representative *f*_3_ statistics for selected population triplets.

**Figure 3.**
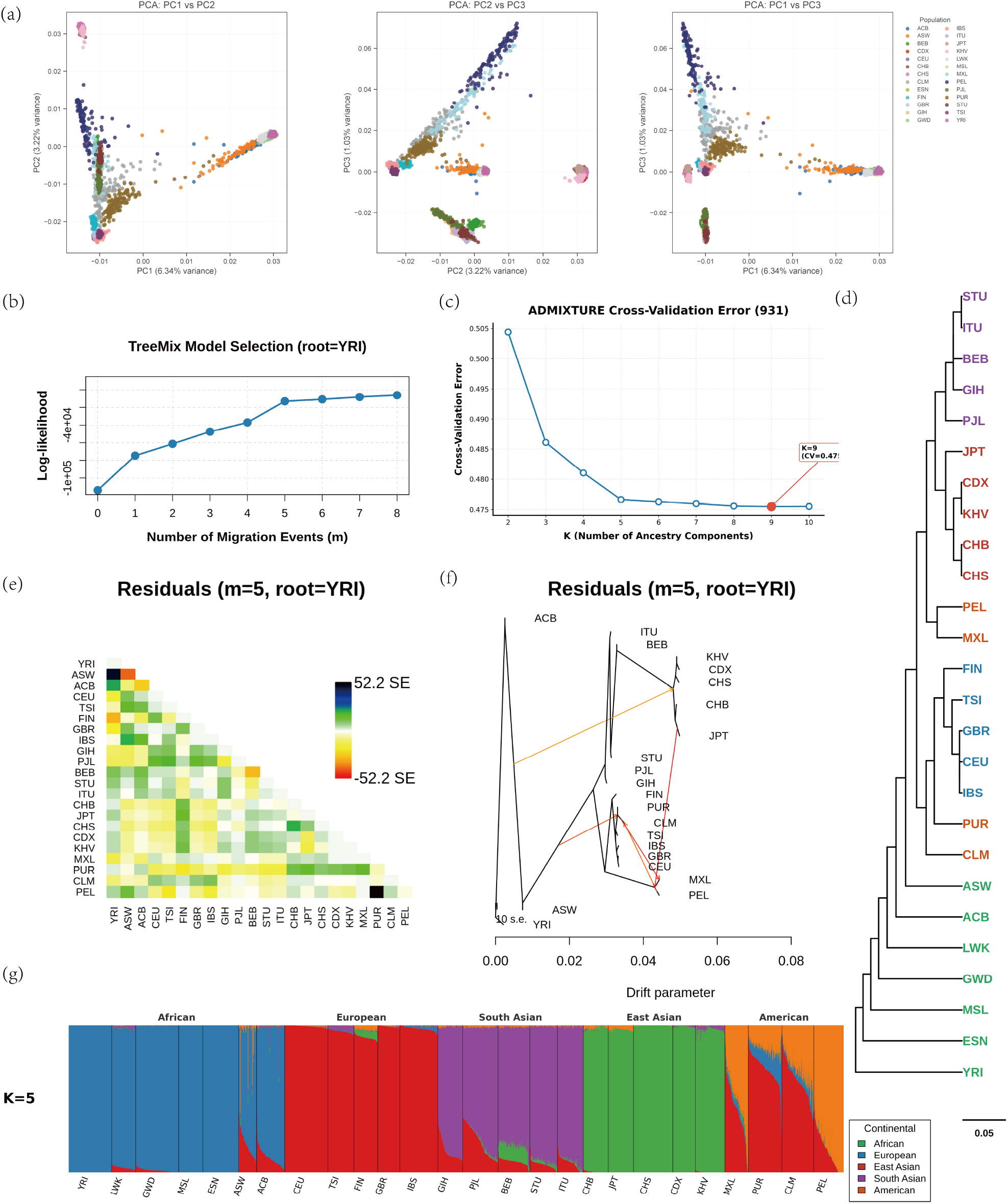
Population-structure and graph-based outputs from the 1000 Genomes example analysis. (a) Principal component analysis (PCA) shown as pairwise projections of the first three principal components with variance explained on each axis. (b) TreeMix log-likelihood as a function of the number of migration edges (*m*) with YRI used as the outgroup for rooting. (c) ADMIXTURE cross-validation error across candidate numbers of clusters (*K*). (d) Population-level clustering dendrogram derived from the mean coordinates of the first five PCs (see Methods). (e) TreeMix residual covariance matrix for the fitted model shown in panel (f). (f) Example fitted TreeMix graph with migration edges. (g) ADMIXTURE ancestry proportions for *K* = 5.

Figure 2 summarizes within-population diversity, autozygosity, and representative allele-sharing output. Panels a and b show observed heterozygosity across populations and its comparison with expected values, and panel c shows the corresponding inbreeding coefficient distributions. Panel d summarizes ROH burden across populations in terms of total ROH length per individual and ROH segment counts. Panel e shows population-specific LD decay curves, and panel f reports representative *f*_3_ statistics with standard errors for selected population triplets.

Figure 3 summarizes complementary structure and graph-based outputs. PCA separates major continental groupings and reveals gradients among populations (Fig. 3a), while a descriptive dendrogram computed from population means of the first five PCs provides a compact population-level summary (Fig. 3d; Methods). ADMIXTURE cross-validation errors are tracked across the tested range of *K* values (Fig. 3c), and Supplementary Fig. S3 shows the corresponding ancestry bar plots. We display the *K* = 5 solution in the main text as a representative continental-scale summary (Fig. 3g).

A series of TreeMix models were generated within the same workflow. Figure 3 shows log-likelihood values across alternative numbers of migration edges (Fig. 3b), residual covariance patterns (Fig. 3e), and fitted graphs (Fig. 3f), while Supplementary Fig. S4 shows additional TreeMix fits and Supplementary Fig. S5 reports the PCA scree plot. Together, these outputs illustrate that PopGenAgent can coordinate Population genetics analysis tools and results within one inspectable workflow context.

## Discussion

Population-genetic analyses are often framed as pipelines, but in practice rely on iterative inspection and revision, where intermediate diagnostics must be kept consistent with downstream results. PopGenAgent addresses this gap by organizing analyses as explicit, revisable plans with preserved intermediate artefacts in a unified context. A key design choice is to constrain execution to curated tools and visualization templates, rather than relying on dynamically generated code, improving consistency and reducing variability across analyses. Compared with existing workflow systems that standardize execution, PopGenAgent focuses on structuring the analysis process itself and maintaining alignment between analytical steps and outputs. Limitations include dependence on predefined components and limited coverage of specialized analyses. Future work will expand supported tools and improve flexibility for more complex workflows.

## Methods

### System Architecture and Persisted Analysis Context

PopGenAgent is a web-based workflow system for Population genetics analyses. In the current implementation, an LLM-based agent serves as a control layer that translates user goals into explicit multi-step plans, retrieves relevant workflow resources, proposes plan revisions in response to intermediate outputs, and assists interpretation and documentation within the current analysis context. Analytical execution is anchored to curated tool and visualization templates rather than free-form tool calling: each step declares required inputs and expected outputs and is instantiated as an inspectable script that calls the underlying analysis program or plotting routine.

Each analysis context is persisted on disk, including the plan, instantiated step scripts, execution logs, and an indexed manifest of generated figures and tables. This persisted state allows users to inspect intermediate artefacts, revise earlier steps, and draft reports without losing the provenance of downstream outputs.

### Workflow Specification Through Plans and Templates

Workflow plans are structured representations of an analysis in which each step is associated with a description, declared input files, and expected outputs (Supplementary Fig. S1). Step templates encode tool-specific command patterns, configurable parameters, and file-organization conventions, whereas visualization templates convert raw outputs into standardized plots and summary panels (Supplementary Fig. S2). Together, the plan and template layers define what the workflow is expected to produce and provide inspectable specifications that can be revised before later steps are executed.

#### Stepwise Execution and Output Validation

Execution proceeds stepwise through the saved plan. After each step, PopGenAgent checks whether the declared outputs are present and satisfy basic integrity criteria, such as non-empty files and expected formats. If a step fails, the system retains diagnostic information for inspection and, when the failure matches a supported template-level adjustment, may attempt a limited retry. This orchestration model is adapted from the framework introduced in BioMaster Su et al. (2025) and specialized here for Population genetics workflows.

#### Implemented Analytical Workflows

The current template library coordinates downstream preprocessing and quality-control tasks (PLINK Chang et al. (2015), VCFtools Danecek et al. (2011), KING Manichaikul et al. (2010)), population-structure analyses (smartPCA/EIGENSOFT Patterson et al. (2006), ADMIXTURE Alexander et al. (2009)), graph-based modeling (TreeMix Pickrell and Pritchard (2012)), and allele-sharing statistics in the *f* -statistics framework (ADMIXTOOLS Patterson et al. (2012)). In the 1000 Genomes example, PopGenAgent starts from a filtered genotype dataset in PLINK format; derived files such as LD-pruned SNP sets and EIGENSTRAT conversions are generated as required by downstream methods.

In addition to tool execution, plotting templates generate standardized panels for ROH and heterozygosity summaries, LD decay curves, PCA projections and scree plots, ADMIXTURE cross-validation curves and ancestry bar plots, and TreeMix likelihood and residual diagnostics (Figures 2–3; Supplementary Figs. S3–S5). For a descriptive population-level summary, we compute Euclidean distances among population means of the first five PCA coordinates and visualize Ward hierarchical clustering as a dendrogram. This visualization is used as a compact summary of population-level separation in PCA space and is not intended as phylogenetic inference.

### Output-Grounded Assistance and Report Generation

Within the current analysis context, PopGenAgent can answer workflow-oriented questions by drawing on recent dialogue, stored planning information, and a curated knowledge store; when configured, it can additionally retrieve relevant PubMed abstracts for background questions. For documentation, the system assembles a Markdown report draft by combining the saved plan with the indexed figures and tables generated during execution, providing a starting point for an analysis write-up grounded in the produced artefacts. Drafted text should be checked against the saved outputs before reuse.

## Data and Code Availability

The analyses reported here use publicly released data from the 1000 Genomes Project, maintained by the International Genome Sample Resource (IGSR). Sequence reads, alignments, and phased variant calls for the project can be accessed through the IGSR portal: internationalgenome.org. An archived release associated with this manuscript is available at zenodo.org/records/18919671.

The PopGenAgent source code is publicly available at github.com/ai4nucleome/POPGENAGENT. The public repository includes an environment.yml file for software setup. Supplementary figures are provided in a separate Supplementary Information file.

## Competing interests

The authors declare no competing interests.

### Generative AI statement

During the preparation of this work, the authors used ChatGPT to assist with language editing and manuscript revision. After using this tool or service, the authors reviewed and edited the content as needed and take full responsibility for the content of the publication.

## References

D. H. Alexander, J. Novembre, and K. Lange. Fast model-based estimation of ancestry in unrelated individuals. Genome research, 19(9):1655–1664, 2009.

A. Auton, G. R. Abecasis, D. M. Altshuler, R. M. Durbin, D. R. Bentley, A. Chakravarti, A. G. Clark, P. Donnelly, E. E. Eichler, P. Flicek, et al. A global reference for human genetic variation. Nature, 526(7571):68–74, 2015.

M. Bourgey, R. Dali, R. Eveleigh, K. C. Chen, L. Letourneau, J. Fillon, M. Michaud, M. Caron, J. Sandoval, F. Lefebvre, et al. Genpipes: an open-source framework for distributed and scalable genomic analyses. Gigascience, 8(6):giz037, 2019.

C. C. Chang, C. C. Chow, L. C. Tellier, S. Vattikuti, S. M. Purcell, and J. J. Lee. Second-generation plink: rising to the challenge of larger and richer datasets. Gigascience, 4(1):7, 2015.

Y. Chang, X. Wang, J. Wang, Y. Wu, L. Yang, K. Zhu, H. Chen, X. Yi, C. Wang, Y. Wang, et al. A survey on evaluation of large language models. ACM Transactions on Intelligent Systems and Technology, 15(3):1–45, 2024.

P. Danecek, A. Auton, G. Abecasis, C. A. Albers, E. Banks, M. A. DePristo, R. E. Handsaker, G. Lunter, G. T. Marth, S. T. Sherry, et al. The variant call format and vcftools. Bioinformatics, 27(15):2156–2158, 2011.

P. A. Ewels, A. Peltzer, S. Fillinger, H. Patel, J. Alneberg, A. Wilm, M. U. Garcia, P. Di Tommaso, and S. Nahnsen. The nf-core framework for community-curated bioinformatics pipelines. Nature biotechnology, 38(3):276–278, 2020.

M. B. Hamilton. Population genetics. John Wiley & Sons, 2021.

K. Huang, S. Zhang, H. Wang, Y. Qu, Y. Lu, Y. Roohani, R. Li, L. Qiu, J. Zhang, Y. Di, et al. Biomni: A general-purpose biomedical ai agent. bioRxiv, page 2025.05.30.656746, 2025. doi: 10.1101/2025.05.30.656746.

Z. N. Kamvar, J. C. Brooks, and N.J. Grünwald. Novel r tools for analysis of genome-wide population genetic data with emphasis on clonality. Frontiers in genetics, 6:208, 2015.

E. Kasneci, K. Seßler, S. Küchemann, M. Bannert, D. Dementieva, F. Fischer, U. Gasser, G. Groh, S. Günnemann, E. Hüllermeier, et al. Chatgpt for good? on opportunities and challenges of large language models for education. Learning and individual differences, 103:102274, 2023.

N. M. Kopelman, J. Mayzel, M. Jakobsson, N. A. Rosenberg, and I. Mayrose. Clumpak: a program for identifying clustering modes and packaging population structure inferences across k. Molecular ecology resources, 15(5):1179–1191, 2015.

J. W. Leigh, D. Bryant, and S. Nakagawa. Popart: full-feature software for haplotype network construction. Methods in Ecology & Evolution, 6(9):1110–1116, 2015.

Y.-L. Li and J.-X. Liu. Structureselector: A web-based software to select and visualize the optimal number of clusters using multiple methods. Molecular ecology resources, 18(1):176– 177, 2018.

A. Manichaikul, J. C. Mychaleckyj, S. S. Rich, K. Daly, M. Sale, and W.-M. Chen. Robust relationship inference in genome-wide association studies. Bioinformatics, 26(22):2867–2873, Journal, 2026, Volume XX, Issue x 7 2010.

N. Park, H. Kim, J. Oh, J. Kim, C. Heo, and J. Kim. Papipe: A pipeline for comprehensive population genetic analysis. Molecular Biology and Evolution, 41(3):msae040, 2024.

N. Patterson, A. L. Price, and D. Reich. Population structure and eigenanalysis. PLoS Genetics, 2(12):e190, 2006. doi: 10.1371/journal.pgen.0020190.

N. Patterson, P. Moorjani, Y. Luo, S. Mallick, N. Rohland, Y. Zhan, T. Genschoreck, T. Webster, and D. Reich. Ancient admixture in human history. Genetics, 192(3):1065–1093, 2012. doi: 10.1534/genetics.112.145037.

J. K. Pickrell and J. K. Pritchard. Inference of population splits and mixtures from genome-wide allele frequency data. PLoS Genetics, 8(11):e1002967, 2012. doi: 10.1371/journal.pgen.1002967.

H. Su, W. Long, and Y. Zhang. Biomaster: Multi-agent system for automated bioinformatics analysis workflow. bioRxiv, page 2025.01.23.634608, 2025. doi: 10.1101/2025.01.23.634608.

A. Webb, J. Knoblauch, N. Sabankar, A. S. Kallur, J. Hey, and A. Sethuraman. The pop-gen pipeline platform: a software platform for population genomic analyses. Molecular Biology and Evolution, 38(8):3478–3485, 2021.

L. Wratten, A. Wilm, and J. Göke. Reproducible, scalable, and shareable analysis pipelines with bioinformatics workflow managers. Nature methods, 18(10):1161–1168, 2021.

W. X. Zhao, K. Zhou, J. Li, T. Tang, X. Wang, Y. Hou, Y. Min, B. Zhang, J. Zhang, Z. Dong, et al. A survey of large language models. arXiv preprint 2303.18223, 2023. doi: 10.48550/arXiv.2303.18223.

J. Zhou, B. Zhang, G. Li, X. Chen, H. Li, X. Xu, S. Chen, W. He, C. Xu, L. Liu, et al. An ai agent for fully automated multi-omic analyses. Advanced Science, 11(44):2407094, 2024.

