## Supplemental Figure S1-S5 for "PopGenAgent: An Agent-Mediated Workflow for Population Genetics Analyses"

### Supplementary Information for: PopGenAgent: An Agent-Mediated Workflow Framework for Population Genetics Analyses

#### Supplementary Figures

##### Workflow State and Stored Resources

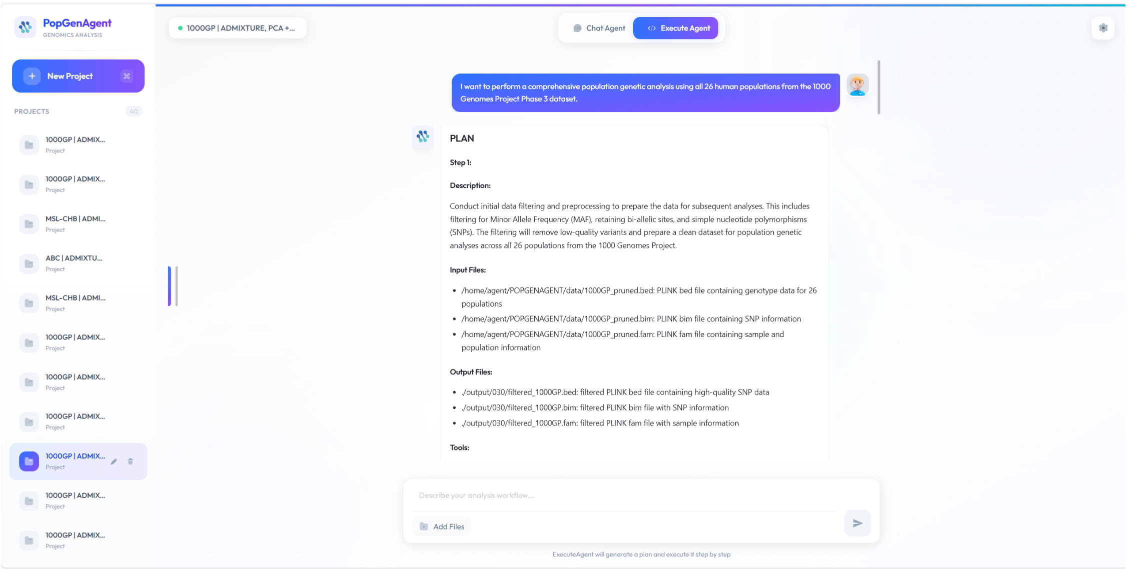


Figure S1: Example plan view from the 1000 Genomes analysis. The panel shows one stored workflow step together with its description, declared input files, and expected outputs.

##### Additional 1000 Genomes Outputs

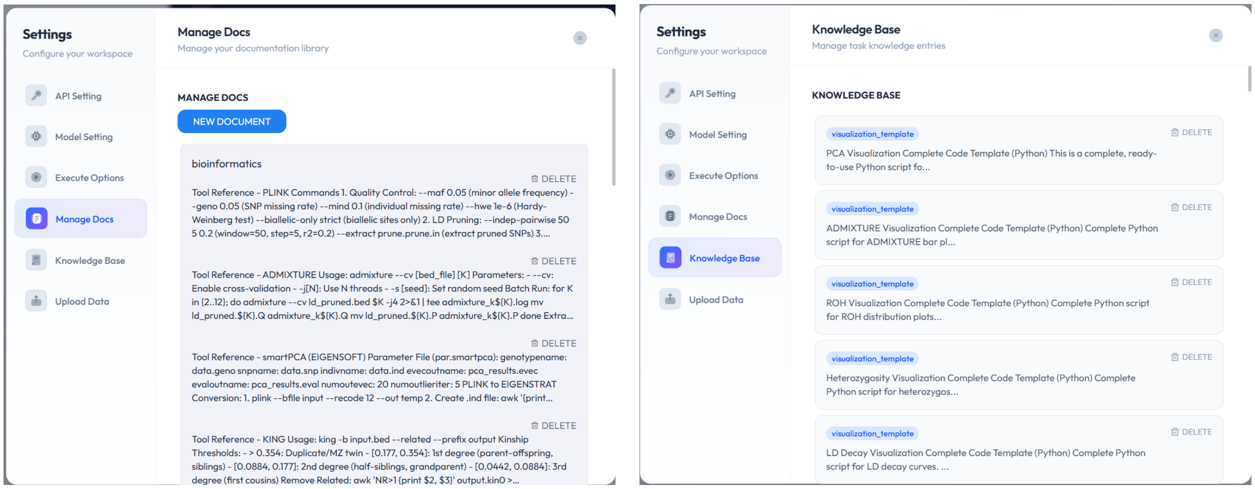


Figure S2: Stored workflow resources used by PopGenAgent. Left: tool-reference records containing command templates and usage notes. Right: reusable task and visualization template entries in the knowledge store.

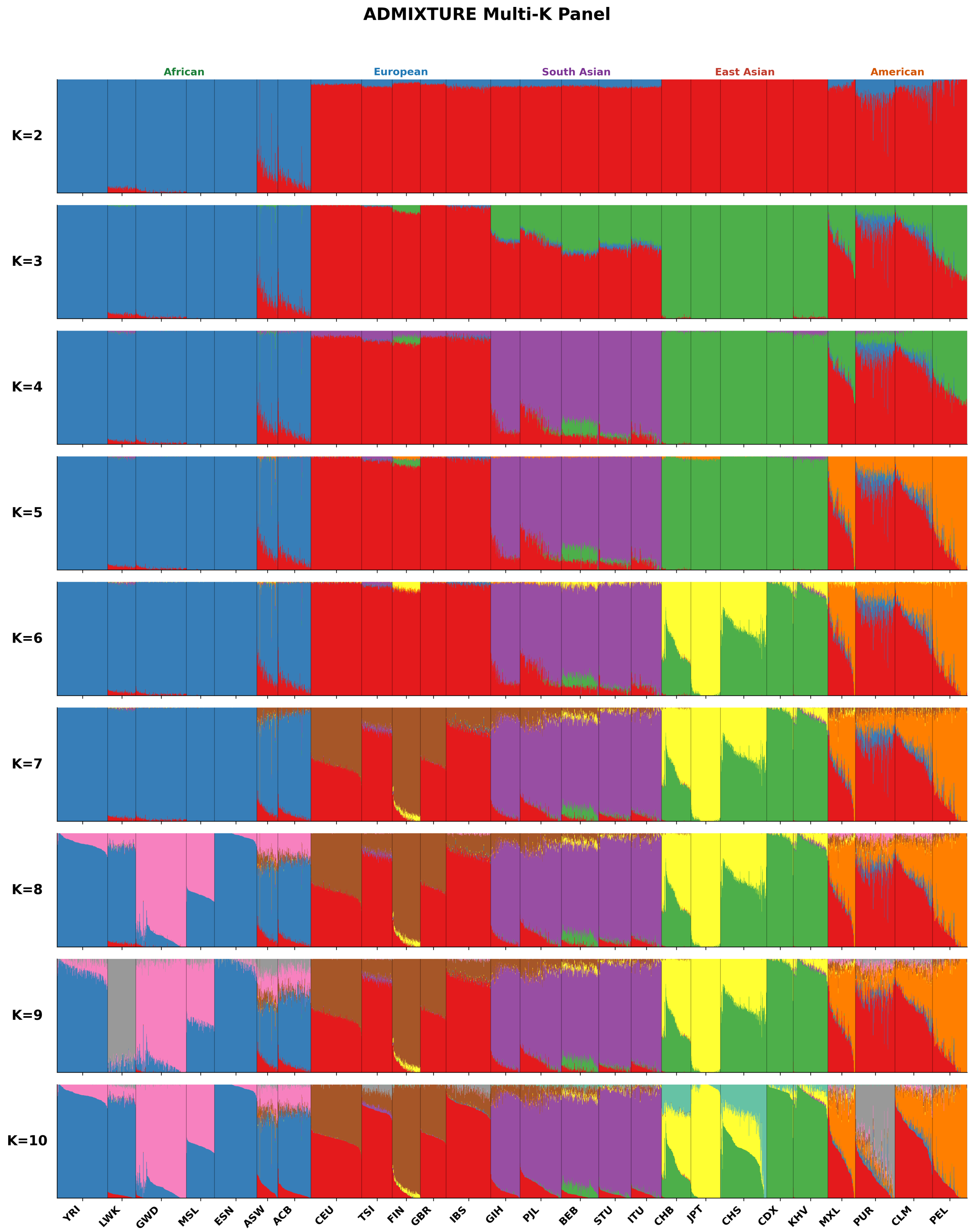


Figure S3: Expanded ADMIXTURE ancestry plots from the 1000 Genomes analysis across the tested range of $K$ values, $K = 2$ to $K = 10$.

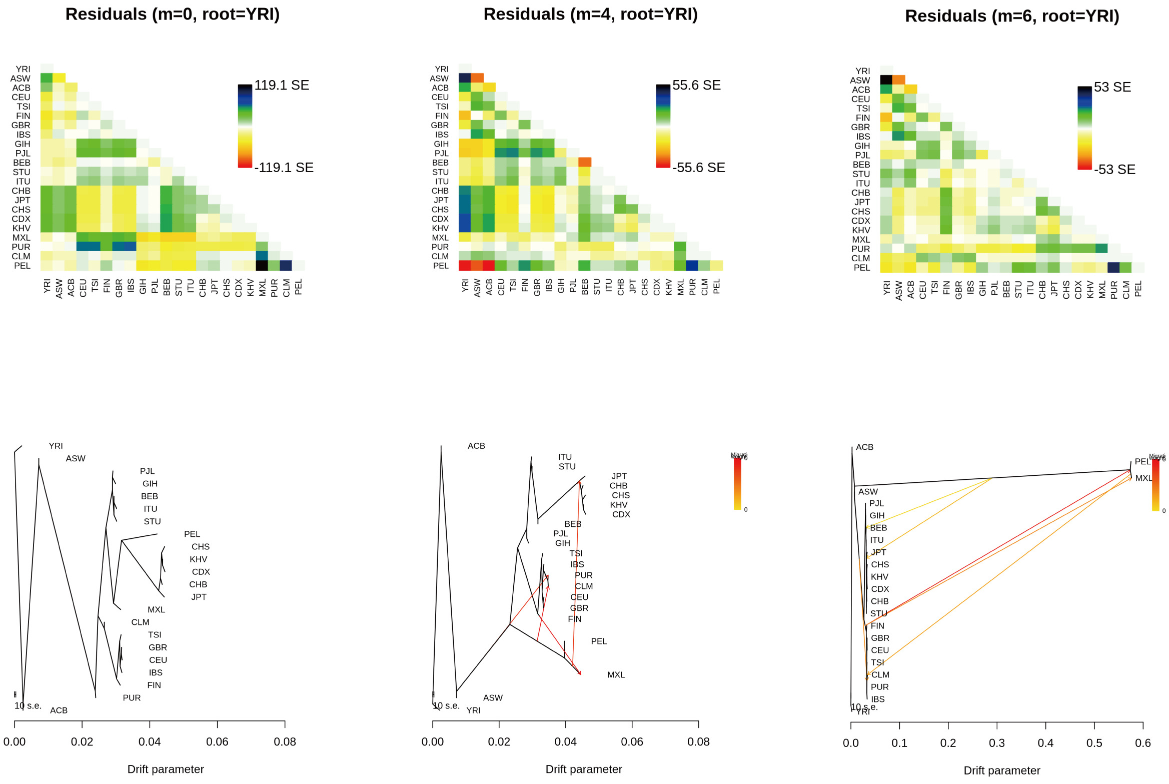


Figure S4: Additional TreeMix fits from the 1000 Genomes analysis. Left: output for $m = 0$, showing the fitted tree together with the corresponding residual covariance panel. Middle: output for $m = 4$, showing a higher-complexity fit from the same scan. Right: output for $m = 6$, showing a higher-complexity fit from the same scan.

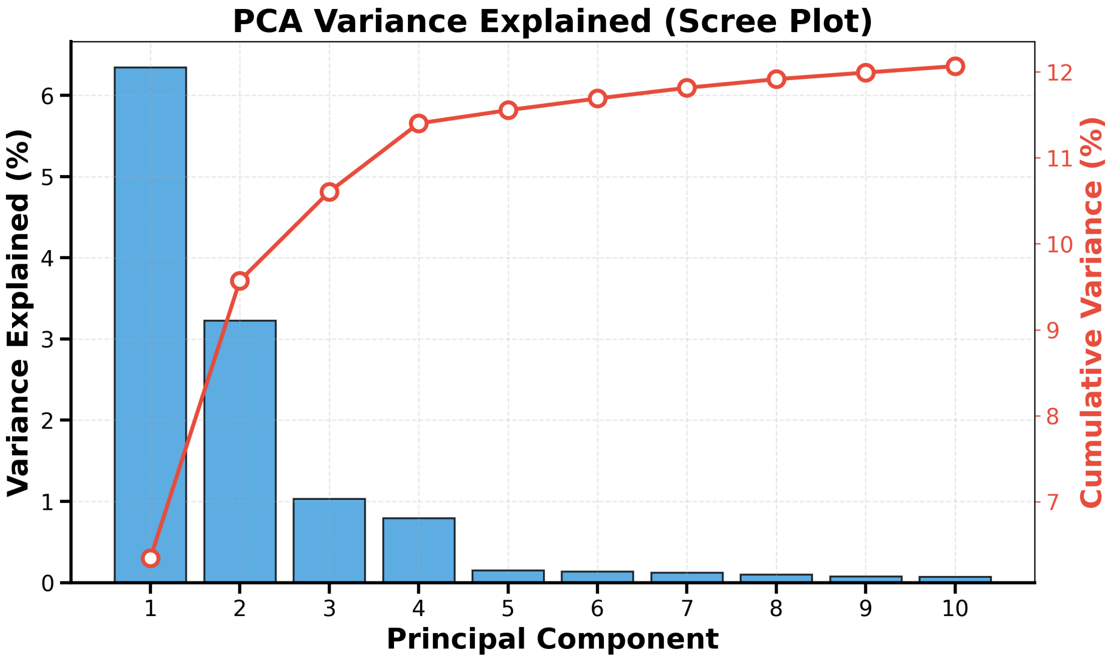


Figure S5: Scree plot for the PCA used in the 1000 Genomes analysis. Bars show the variance explained by individual principal components, and the line shows cumulative variance across components.
